# Assessing predictors for new post translational modification sites: a case study on hydroxylation

**DOI:** 10.1101/2020.02.17.952127

**Authors:** Damiano Piovesan, Andras Hatos, Giovanni Minervini, Federica Quaglia, Alexander Miguel Monzon, Silvio C.E. Tosatto

## Abstract

Post-translational modification (PTM) sites have become popular for predictor development. However, with the exception of phosphorylation and a handful of other examples, PTMs suffer from a limited number of available training examples and their sparsity in protein sequences. Here, proline hydroxylation is taken as an example to compare different methods and evaluate their performance on new experimentally determined sites. As a proxy for an effective experimental design, predictors require both high specificity and sensitivity. However, the self-reported performance is often not indicative of prediction quality and detection of new sites is not guaranteed. We have benchmarked seven published hydroxylation site predictors on two newly constructed independent datasets. The self-reported performance widely overestimates the real accuracy measured on independent datasets. No predictor performs better than random on new examples, indicating the refined models are not sufficiently general to detect new sites. The number of false positives is high and precision low, in particular for non-collagen proteins whose motifs are not conserved. In short, existing predictors for hydroxylation sites do not appear to generalize to new data. Caution is advised when dealing with PTM predictors in the absence of independent evaluations, in particular for unique specific sites such as those involved in signalling.

**Author Summary:** Machine learning methods are extensively used by biologists to design and interpret experiments. Predictors which take the only sequence as input are of particular interest due to the large amount of sequence data available and self-reported performance is often very high. In this work, we evaluated post-translational modification (PTM) predictors for hydroxylation sites and found that they perform no better than random, in strong contrast to performances reported in the original publications. PTMs are chemical amino acids alterations providing the cell with conditional mechanisms to fine tune protein function, thereby regulating complex biological processes such as signalling and cell cycle. Hydroxylation sites are a good PTM test case due to the availability of a range of predictors and an abundance of newly experimentally detected modification sites. Poor performances in our results highlight the overlooked problem of predicting PTMs when best practices are not followed and training data are likely incomplete. Experimentalists should be careful when using PTM predictors blindly and more independent assessments are needed to separate the wheat from the chaff in the field.

## Introduction

Post translational modifications (PTMs) are alterations of the primary structure of the protein. PTMs include both new covalent links and cleavage events. Almost every protein in the cell undergoes modification during its lifetime (1). More than 600 different amino acid modifications are catalogued in UniProtKB (2). PTMs provide a way to expand the spectrum of protein functions and an additional layer for pathway regulation (3). They are catalyzed by enzymes that identify a specific site in the substrate protein. A plurality of PTM motifs reside in intrinsically disordered regions in order to provide enzyme accessibility (4). Over the last few years, a deluge of methods have been proposed to predict PTM sites from sequence. For a recent review see e.g. (5). The reasons for this popularity are broadly twofold. Given the paucity of experimental data and relevance of PTMs for cellular regulation, there is a legitimate expectation that computational tools should fill in the experimental void. Computational tools can become fundamental as hypothesis generators for an effective design of PTM experiments. The implementation of predictors is straightforward thanks to the sequence specificity and peculiar physico-chemical properties of PTM motifs. This simplicity makes PTM prediction from sequence easily accessible to machine learning methods, but also presents several potential pitfalls (6). In order to be useful for experimentalists, PTM predictors should provide good performance and be robust. Performance should be high enough to limit false positives to a minimum, while ensuring sufficient coverage. Perhaps more importantly, the method should be robust enough to maintain performance across a range of different datasets since it is often not clear what experimental conditions may introduce biases. On both accounts, PTM predictors may be problematic as they are rarely assessed by independent third parties. Indeed, their ability to identify new modification sites has been questioned (7) and effective results have been obtained only for a few PTM types (5). The problem of validating machine learning methods has already been raised and best practices have been proposed (6). The self-reported accuracy may be overestimated and PTM predictors do not perform better than random when adopting the wrong training strategy, leading to overfitting (7). Generalizing models for PTM site recognition is difficult. The number of experimental observations is low, with a lot of new types of motifs not yet available.

In this work, proline hydroxylation is taken as a case study to answer the question how useful PTM predictors, especially those trained on small datasets, are to design experiments. Hydroxylation is one of the most abundant PTMs in the cell (8). However, despite improvements in mass-spectrometry (MS) techniques, likely only a small fraction of all hydroxylated sites has been experimentally detected so far. Proline hydroxylation (PH) is a PTM carried out by prolyl hydroxylases catalyzing the addition of an hydroxyl group to the sidechain pyrrolidine ring at the gamma position. This modification is crucial for correct folding of the collagen triple-helix, which contains the conserved xPG motif. PH also plays a crucial role in signaling, in particular in the oxygen sensing pathways, including angiogenesis (9) and tumor cell proliferation (10, 11). An example is HIF1α, the main target of the von Hippel-Lindau (pVHL) E3 ubiquitin ligase complex (12). In normoxia, the prolyl hydroxylase domain-containing enzymes (PHDs) hydroxylate HIF1α, promoting its degradation through pVHL binding (13). Under low oxygen concentration, the PHDs are inactivated and HIF-1α translocates into the nucleus to activate vascular proliferation and neoangiogenesis genes (14).

The first hydroxylation predictor (15) was trained to predict only collagen modifications. Several predictors exist as web servers: HydPred (16), PredHydroxy (17), RF-Hydroxysite (18), iHyd-PseAAC (19) and iHyd-PseCp (20). The latter has not been considered in our analysis as the server is unstable and freezes frequently. The stand-alone software OH-Pred (21), ModPred (4) and AMS3 (1) are also available. All are potential tools for large-scale analysis, taking only the protein sequence as input. Implementations include standard machine learning algorithms like Support Vector Machines (SVM), artificial Neural Networks (NN) and Random Forests (RF), as well as alternative techniques like logistic regression and probabilistic classifiers. All methods were trained on SwissProt (22) annotation but with different strategies to define positive and negative examples and different approaches to evaluate model quality. None of them used a real independent dataset for validation, i.e. unaffected from SwissProt biases.

Here, we evaluate methods considering separately collagen and signalling examples as well as single proteins versus high throughput MS experiments. The majority of new hydroxylated prolines (Hyp) come from two MS experiments, one on HeLa cells and another based on a large experiment made on multiple tissues and samples, recently published and not yet available in public databases when predictors were trained (23–25). The number of MS hydroxylated sites is comparable to the entire SwissProt database. In this work, we considered new data extracted from two different high-throughput MS experiments to compare different predictors and evaluate their ability to generalize for new (unseen) motifs. The new example sources allowed us to perform a real unbiased blind test. A Naïve HMM predictor trained including MS data, has been implemented to simulate the effect of integrating new examples. The analysis presented here provides a starting point for a critical discussion on the problem of reliably predicting new PTMs.

## Results

PTM experiments are usually designed with limited time and budget. Prediction tools have the potential to make experiments more effective. However, effective predictors need to work much better than random. In particular, in the case of PTMs, where the fraction of modified residues is low, the false positive rate (fraction of false positives over negative examples) should be minimized. Due to the low coverage of PTM evidence in public databases, predictors should be able to generalize, i.e. identify new motifs never seen before. In the following, different predictors are evaluated against new hydroxylation sites and various problems related to PTM prediction are discussed. In order to provide an objective evaluation we considered “old” examples (Literature), i.e. those available from the SwissProt database at the training time, “new” examples coming from two different mass-spectrometry experiments (MS-Kim, MS-HeLa) and “new but easy” examples (MS-collagen) which are a subset of the “new” examples found in collagen proteins and therefore similar to the “old” well known collagen motifs. Table 1 shows the main splits used in the paper. For methods which provide a confidence score, the Precision-recall and ROC curves are reported in Supplementary Figures S3-S8.

**Table 1.** Datasets. Negative clusters (“Negative”) contain only clusters with non-hydroxylated sites. Others datasets have clusters with both positive and negative examples, but negatives are completely removed during evaluation (*). Negative sites (^) considered during evaluation are always resampled for each replica, based on the size of the positive dataset.

| Dataset | Clusters | Evaluated sites | Filtered negative sites* |
| --- | --- | --- | --- |
| Collagen | 5 | 243 | 4,517 |
| Literature-collagen | 4 | 152 | 4,588 |
| MS-collagen | 4 | 95 | 4,508 |
| Literature | 167 | 877 | 13,481 |
| MS-HeLa | 198 | 324 | 14,982 |
| MS-Kim | 625 | 1,694 | 24,692 |
| MS | 705 | 2,002 | 26,631 |
| Negative | 493 | 7,875<sup>^</sup> | - |

### Predictor performance

As a starting point, Table 2 shows details about the evaluated predictors, including self-reported performance. Self-reported performance is taken from the corresponding publications selecting, when possible, values calculated on independent validation sets, i.e. excluding training examples.

**Table 2.**
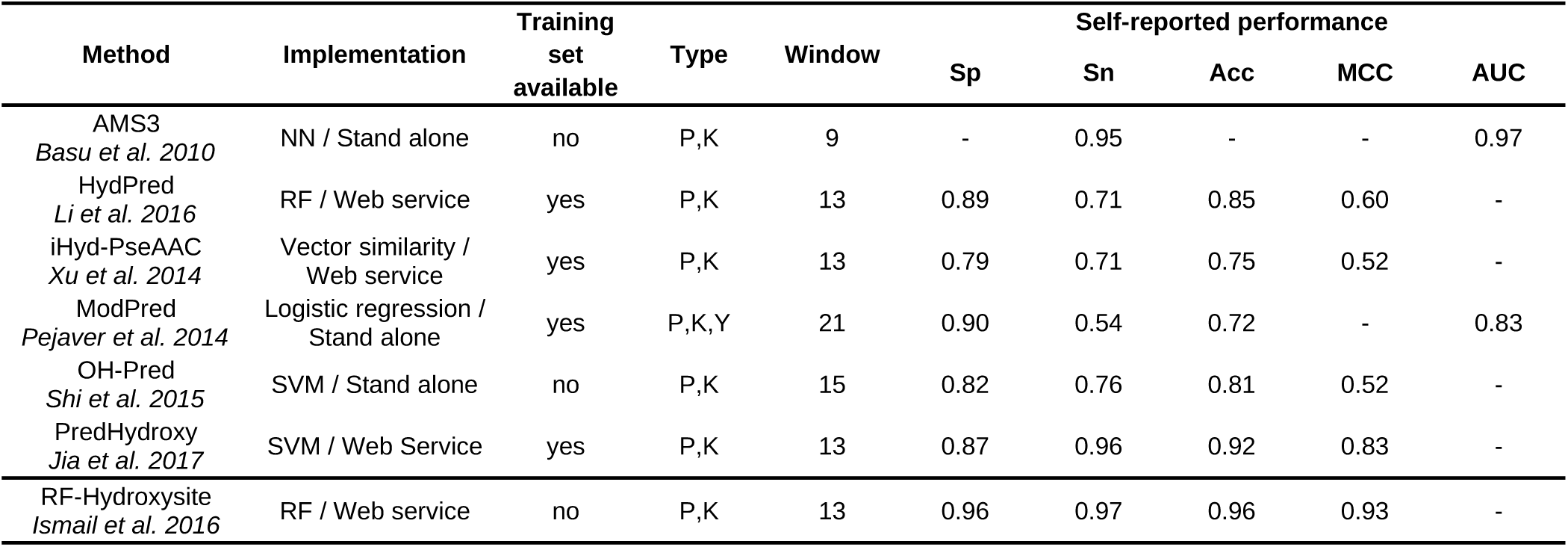
Methods overview. Self-reported performance is taken from the corresponding method publications preferring values reported from independent validation sets, i.e. not used in the training.

In Figure 1 the performance considering manually curated example from single protein experiments (Literature) is shown to simulate the evaluation provided by the method publications. The majority of “Literature” examples in fact come from SwissProt and were already available at the training time. ModPred, HydPred and OH-Pred perform as declared. Instead, ASM3, PredHydroxy and RF-Hydroxysite show a decrease, with the latter performing worse than random and with a negative MCC. The RF-Hydroxysite web server probably suffers a software bug.

**Figure 1.**
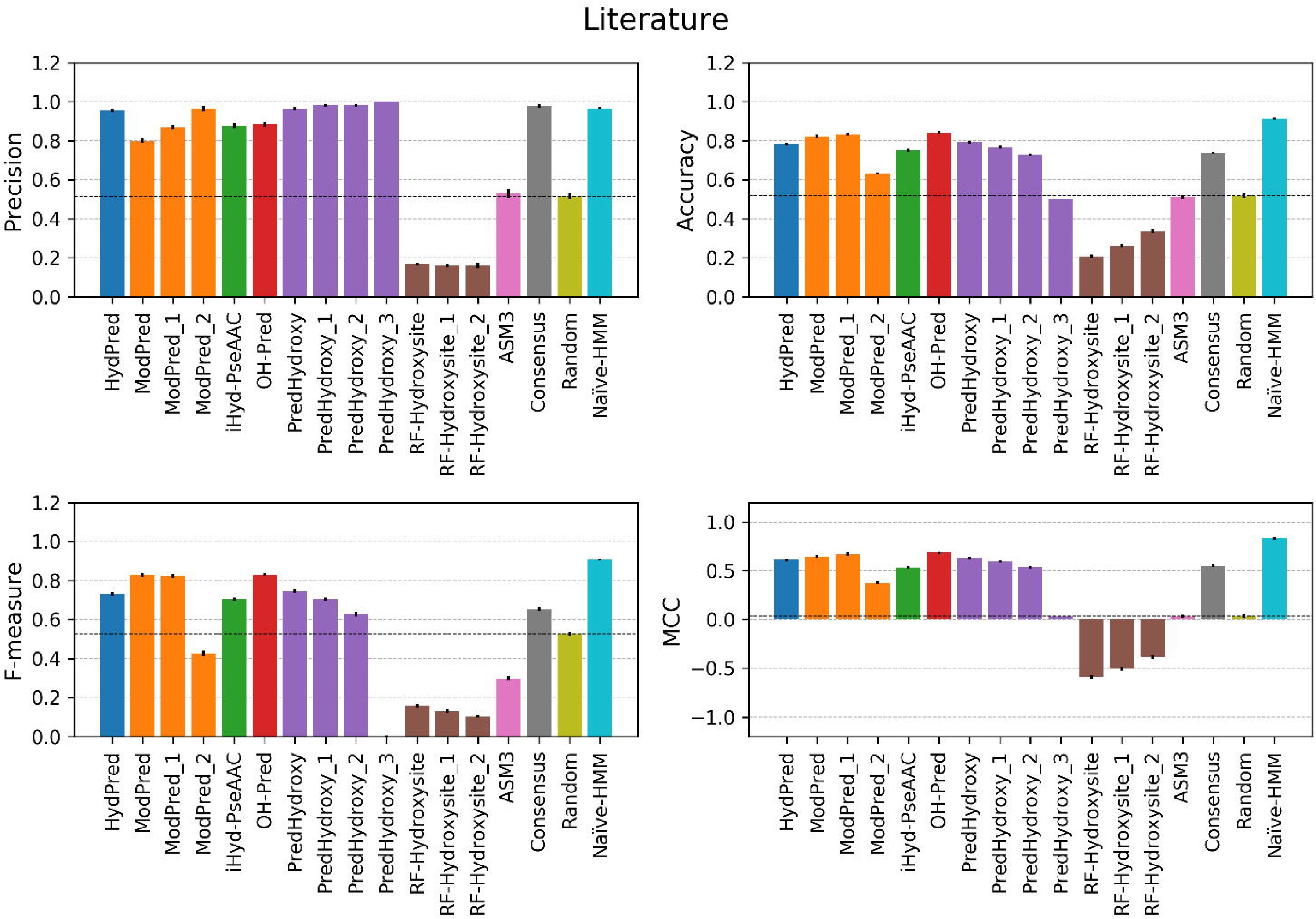
Performance on literature examples. The evaluation is performed only considering hydroxylated sites detected by single protein experiments (Literature dataset). Error bars are the standard deviation calculated over 1,000 replica sets. The consensus baseline method is the majority vote across all predictors. Suffix numbers in the method names indicate increasing quality threshold as defined by developers.

For best methods, sensitivity and specificity are both high (Supplementary Table 1). Methods providing a confidence threshold can modulate the precision (PPV) correctly, with the exception of PredHydroxy for which, at high confidence (0.9) the sensitivity goes to zero and behaves like a random predictor.

Both absolute values and predictor rankings change significantly when measuring the performance on new MS examples (Supplementary Table S4). Even if there are substantial differences in the MS site detection protocol between the MS-HeLa and MS-Kim datasets, predictors behavior is very similar (Figure 2, Supplementary Tables S2, S3). All methods do not seem to generalize well and have a low sensitivity. High specificity combined with low sensitivity, as for PredHydroxy, is critical in particular for unbalanced and incomplete datasets like PTMs. For Hyp the positive to negative ratio in the human genome is less than 0.10 and negative examples might become positive as new experimental evidence is collected. All predictors have a balanced accuracy close to 0.5, indicating a random behavior. Notably, only ModePred is better than random for the Ms-HeLa dataset (Figure 2, Supplementary Table S2), it achieves the best MCC (0.13 MS-Kim, 0.32 Ms-HeLa), but still not sufficient for practical use by experimentalists. For example, considering the merged MS dataset, only 62% of hydroxylation sites predictions will be correct (precision) and 65% of modified residues undetected (false negative rate) (supplementary Table S4). We explored the possibility of reducing false positives by implementing a consensus predictor based on a majority vote. In all evaluations the consensus is in line with methods average. Overall, it is fair to say that the predictors do not work on the new datasets. The NaÏve-HMM baseline (see Material & Methods), which is trained considering also new examples, instead behaves like a perfect predictor. While NaÏve-HMM cannot be considered free from bias as its validation set overlaps the training set, it demonstrates how negative sites are significantly different from positives and predictors can benefit from using them in the training.

**Figure 2.**
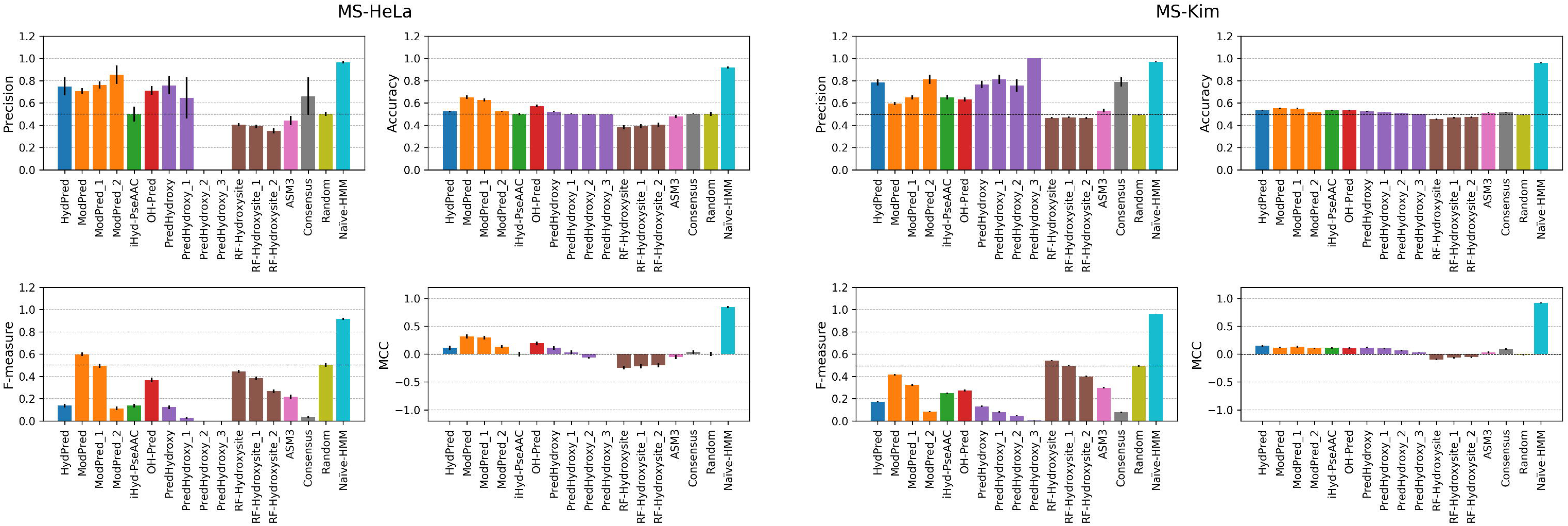
Performance on the MS-HeLa and MS-Kim datasets. The evaluation is performed only considering hydroxylated sites detected by mass-spectrometry experiments (MS-HeLa and MS-Kim dataset). Consensus and errors are calculated as in the previous figure. Suffix numbers in the method names indicate increasing quality threshold as defined by developers.

### Collagen

A case of special interest may be collagen, which accounts for very specific hydroxylation motifs. In collagen, hydroxylation affects different locations corresponding to different sites and molecular meanings. Collagen presents the canonical XaaYaaGly pattern with the ProHypGly triplet found in 10.5% of collagen motifs (37). Xaa is a Pro in 28% and Yaa is Hyp in 38% of the cases. Hyp position matters as in Xaa it prevents the formation of the tropocollagen (TC) triple helix (38). Collagen motifs are conserved, well studied and therefore easier to predict compared to signaling hydroxylation. Both the Literature and MS datasets include 152 and 95 collagen sites respectively (Table 1), all grouped into only 5 different clusters (Supplementary Figure S2). Unsurprisingly, considering Literature collagen motifs, predictors stand out achieving a maximum MCC of 0.81 and accuracy of 0.94 (Supplementary Table S3). The situation is similar when considering collagen examples from the MS dataset (Figure 3) indicating the quality of the MS data is comparable to Literature.

**Figure 3.**
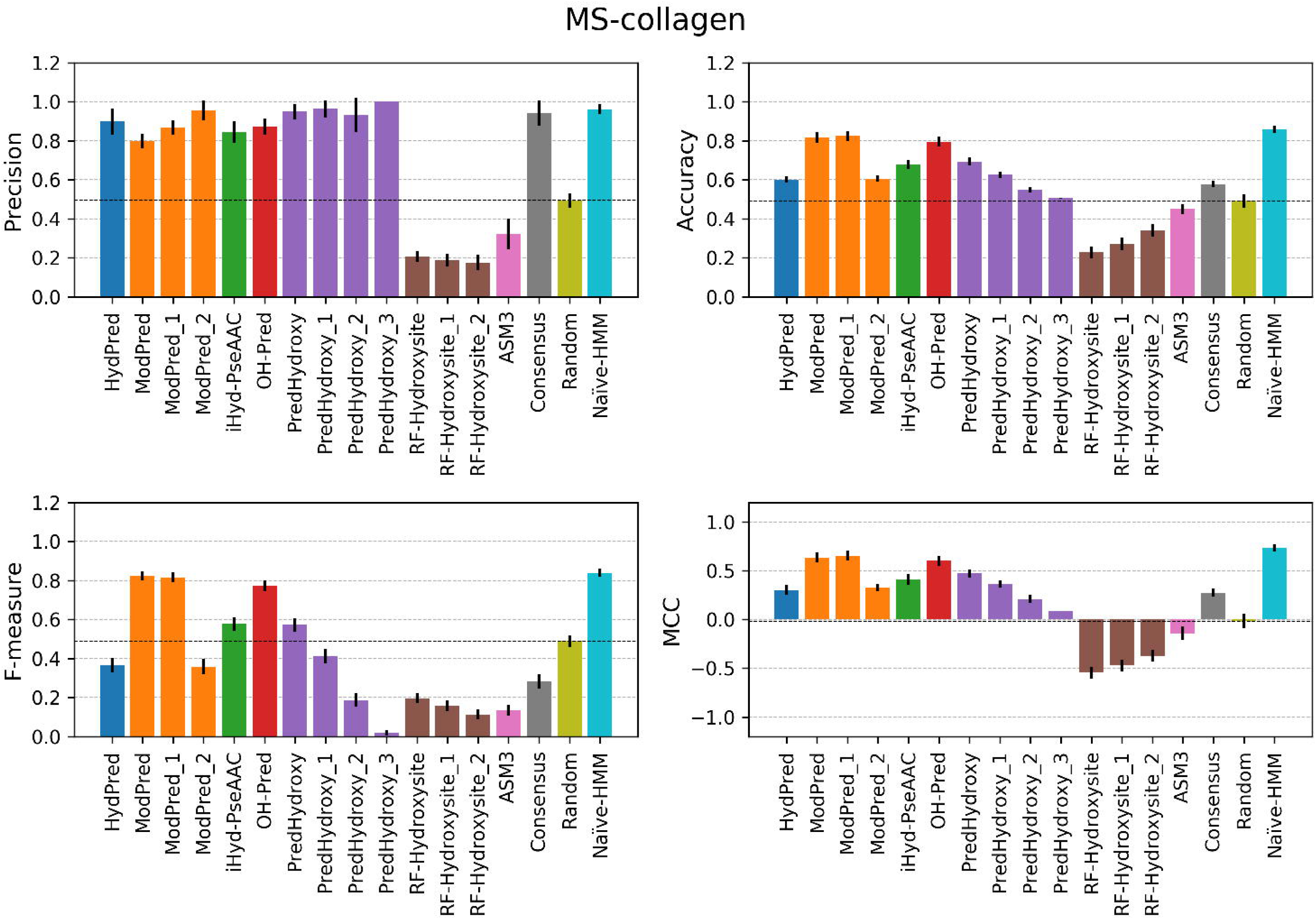
Performance on MS-collagen examples. The evaluation is performed only considering hydroxylated sites detected by mass-spectrometry experiments and belonging to collagen proteins (MS-collagen dataset). Consensus and errors are calculated as in the previous figure. Suffix numbers in the method names indicate increasing quality threshold as defined by developers.

### Dataset characterization

Hydroxylation is known to be linked to angiogenesis and tumor growth and hydroxylases have been observed to be particularly active (39) with changing collagen patterns in cancer cells (40). Therefore, we distinguished between examples from single-protein experiments described in the literature and hydroxylation observed in tumor cells from MS experiments. MS experiments are not free from bias (41) and are enriched in flexible peptides (42). Also, one of the two MS experiments has been performed using anti-hydroxyproline antibody which may include a sequence bias. In this study we limited the analysis to high confidence sites with at least 80% experimental score probability.

Considering site sequence similarity the Literature dataset is a subset of the MS dataset. 122 out of 165 Literature clusters (74%) have at least one MS site (intersection). These clusters include 92% of the total Literature examples (13,196 sites). On the other end, 583 clusters including 15,437 sites have only MS examples representing the real new hydroxylation motifs.

In order to assess non-specific proline binding, a comparison between MS and Literature datasets is reported in Figure 4. The residue frequencies around the modified proline (Panel A) decay exponentially for the MS dataset while Literature sites have a peak at 25% with a distribution shifted toward enriched sites, which is probably due to a stronger contribution of highly repeated collagen patterns. On the other end a supposed bias towards polyproline detection by MS experiments is not observed.

**Figure 4.**
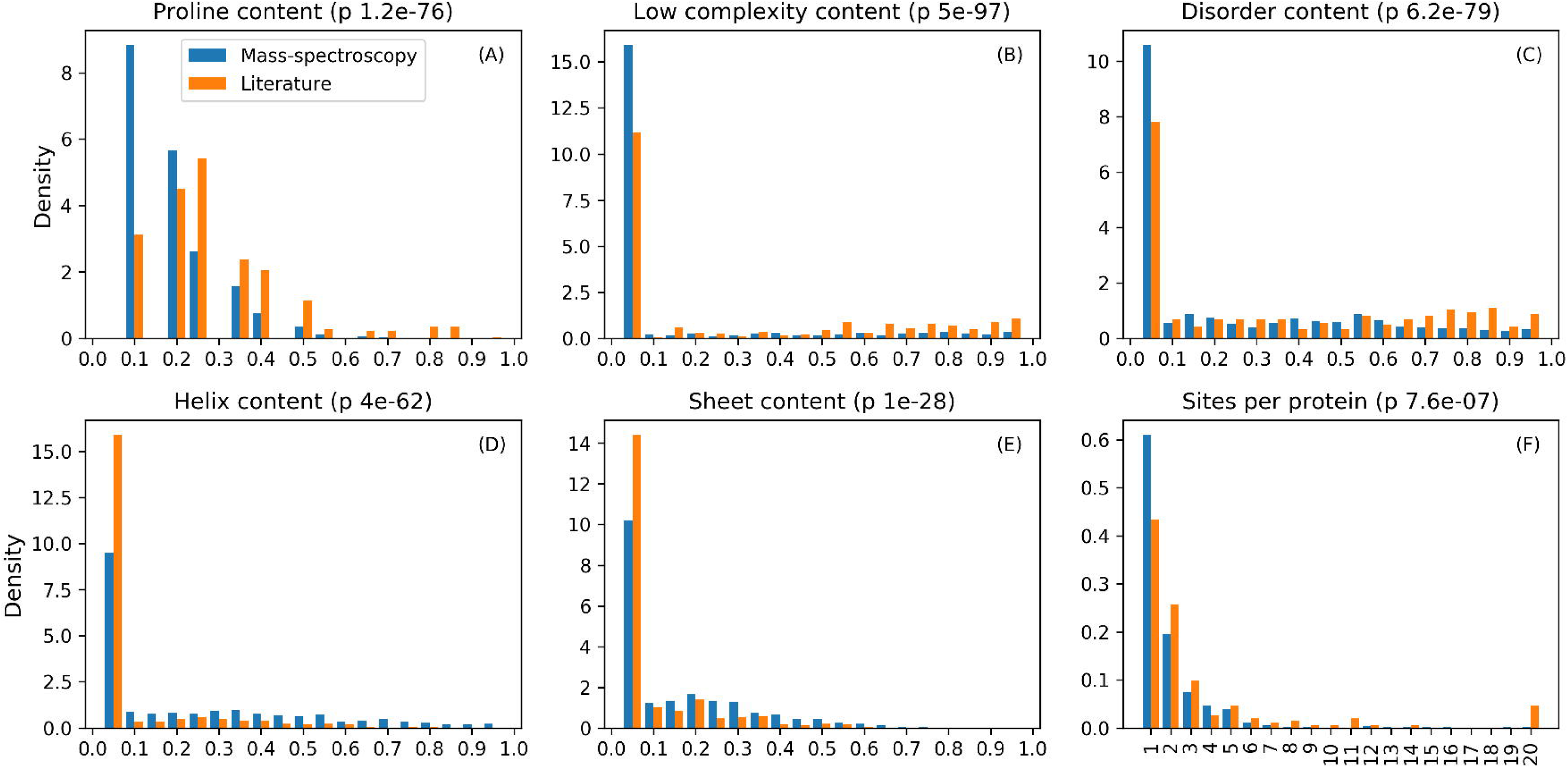
Features distribution for MS and Literature sites. Content refers to the fraction of residues in the site sequence associated with a given feature. Density refers to the fraction of proteins in the dataset with a given number of sites.

The number of hydroxylated sites per protein (Panel F), despite it is similar for the two datasets, shows that Literature is shifted having more sites per proteins, which might also be related to collagen abundance. Again, the expected over-hydroxylation of MS sites is not observed. According to the MS experiment results three quarters of the sites are hydroxylated 100% of the time, but we do not have this information for the Literature dataset. As previously observed for several PTM types (4), hydroxylation also has a preference for intrinsically disordered regions (Panel C) rather than for secondary structure elements (Panel D and E) or low complexity regions (Panel B). In summary, even if the distributions are statistically different, no particular evidence was found for a specific sequence-based bias in the MS dataset compared to Literature examples. This suggests that the predictors, if properly trained, should be able to generalize sufficiently to predict MS hydroxylation sites.

## Discussion

We assessed hydroxylation sites predictors as a paradigm for common situations arising with sequence-based machine learning methods (6). Analyses using unbiased testing data set suggest that predictors perform no better than random when predicting hydroxylation PTM on new examples, which make them unsuitable for experimental biologists. This is in strong contrast to the self-reported performances on independent datasets. Bad performance on new examples can be explained by one of two reasons. First, a bad training protocol and second, the intrinsic problem of machine learning methods that are able to detect only those patterns similar enough to training examples. For the hydroxylation predictors assessed here, problems in the training protocol extend also to the construction of the dataset. Sometimes negatives are chosen from complement sites in hydroxylated sequences, some other times negatives are just randomly selected from non hydroxylated sequences. The first strategy might be more reliable since presumably both positive and negative sites have been tested experimentally, on the contrary, randomly selected proteins might include modifications not observed yet. Some methods generate training datasets filtering negative sites exploiting a third-party solvent accessibility predictor in order to exclude those in the protein surface. This is problematic since it can introduce additional uncertainty when surface residues are mispredicted. Another critical point is the sequence redundancy in the training set. All methods, with the exception of ModPred, reduce redundancy at the protein level. This is problematic since protein pairs with low global identity can share short regions with high similarity including the hydroxylation sites. Even more problematic is the choice of the validation set. When both the training and validation sets include the same bias, predictors will over-weight biased features and perform poorly on new examples. Aside from technical problems related to machine learning, predicting PTMs is particularly difficult because different modification patterns can be observed for different cells and in response to environmental conditions including diseases. Apparently hydroxylation does not escape this paradigm and predictors are not able to provide a ground for generating new hypotheses. New data generated by MS experiments will improve predictor accuracy and sensitivity but at the moment it is hard to estimate the amount of examples necessary to represent the entire PTM motifs space. This is particularly critical for PTMs in general as they are heavily involved in the regulation of biological processes/signaling and have an extremely dynamic turnover.

In conclusion, we have provided a thorough independent assessment of previously published hydroxylation site predictors. Our results do not bode well for the field, suggesting that self-reported performance is often overestimated and difficult to replicate. This should be seen as an example for the common pitfalls associated with many of the current PTM predictors. Knowing how likely training sets cover the real world is crucial. Experimentalists should be careful when using PTM predictors until more independent assessments are able to separate the wheat from the chaff in the field.

## Materials & Methods

### 2.1 Dataset

Hydroxylated substrates were retrieved from SwissProt (26) (version 2018_03) considering all organisms. The dataset is further filtered by retaining only manually curated annotations with evidence code “experimental evidence used in manual assertion” (ECO:0000269) or “curator inference used in manual assertion” (ECO:0000305). UniProtKB provides a controlled vocabulary of all PTM types of which the following terms are considered: 4-hydroxyproline (1,033 sites), hydroxyproline (220 sites), 3-hydroxyproline (27 sites), 3,4-dihydroxyproline (5 sites) and (3R,4R)-3,4-dihydroxyproline (1 site).

Additional sites are retrieved from the literature (27–30) including two large scale MS experiments, one on HeLa cells (23) and another based on 30 normal human samples including almost all tissues (24), reanalysed with a new software, TagGraph (PRIDE accession PXD005912) (25). MS experiments provide the majority of new examples which are currently not included in SwissProt. In order to minimize assignment errors HeLa examples are filtered to retain only sites with a confidence probability of 0.8. Compared to the original analysis in (24), TagGraph on average tripled the number of identified sites with a degree of variability depending on the type of tissue.

Even though some methods predict lysine and tyrosine hydroxylation, only prolines are considered in the assessment in order to reach statistical significance given the paucity of data for other PTMs. All predictors identify modified residues exploiting the sequence context (surrounding residues) with the assumption that it encodes sufficient information for molecular recognition. The maximum window size adopted by the methods used in this study is 21 residues.

The final dataset includes 1,419 proteins which contain at least one hydroxylated proline and for which all predictors give an output. Only 10% of all prolines, 3,771 out of 37,670, are hydroxylated. Sites are defined considering a window of 13 residues centered on the proline. The prediction evaluation has been performed on a subset of sites selected as described below and shown in Figure 5.

**Figure 5.**
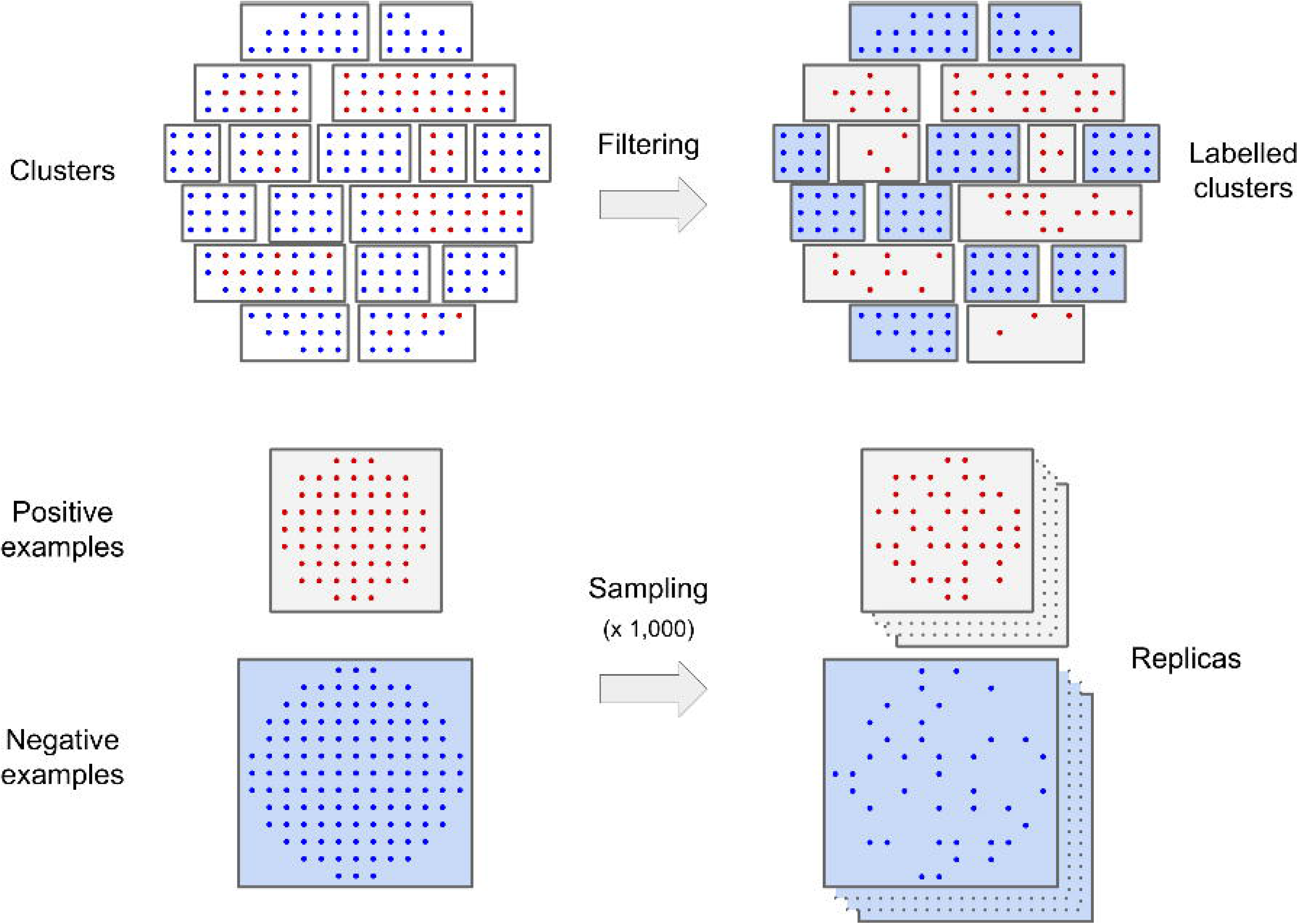
Dataset generation. Negative (blue dots) and positive sites (red dots) are clustered based on sequence similarity. Positive clusters (gray background) contains at least one hydroxylates site and negative examples falling inside positive clusters are removed. 1,000 replica sets are created by random sampling 70% of the positive sites and the same number from the negatives.

Both positive and negative sites are clustered together based on sequence similarity. Clusters are calculated from a distance matrix representing the pairwise sequence divergence between all sites. The distance is computed as the inverse of the similarity score. Similarity is calculated elementwise for each pair of residues of the two sequence sites. The score of a single pair is taken from the Blosum62 matrix, with a penalty of −5 for gap opening and −1 for gap extension. Gaps are introduced to substitute non-canonical residues or to pad the site sequence to reach the window size when a proline is too close to sequence ends. In Supplementary Figure S1 the dendrogram of hierarchical clustering calculated with UPGMA algorithm as implemented in the SciPy Python library is reported. Negative examples too similar to positive examples, i.e. falling inside clusters containing at least one hydroxylated proline (positive site), are removed. The evaluation of the predictors is performed on 1,000 different replica sets. Each replica is built by random picking 70% of positive sites available and the same number of negatives, so to obtain a balanced dataset.

The evaluation is provided for different subsets of positive examples. Namely, sites from single protein experiments (Literature) and mass-spectrometry (MS-Kim, MS-HeLa). The two mass-spectrometry datasets were also merged for some of the presented analysis (MS dataset). Both the Literature and MS datasets can be further divided into collagen and non-collagen entries by recognizing the collagen motif from Pfam domain annotation PF01391 (31). The dataset of collagen sites from single protein experiments is named Literature-collagen and collagen sites from MS is MS-collagen. For each subset and each replica, the tested examples are resampled as described above.

To characterize the dataset sequences, secondary structure (helix/sheet propensity) is predicted using FELLS (32), functional disorder with MobiDB-lite (33) and low complexity with SEG (34). All predictors were executed on the full protein sequence. The fraction of residues assigned to a given feature, i.e. “content”, is calculated for each site. Proline content is calculated as the fraction of prolines in the site irrespective of any hydroxylation modification.

### 2.3 Prediction

We evaluated seven different predictors on entire protein sequences. Some are implemented as stand-alone software and others as web servers. None of them provide any APIs for programmatic access and predictions were taken from web page results. The iHydPse-CP web server (20) stopped working during the benchmark and was excluded from the evaluation as it was never restored. Another method is described in the literature (35) but the software has not been released even upon request. Some methods are designed to predict different modification types. Our evaluation focuses only on proline hydroxylation in order to provide statistically significant results. Some predictors (HydPred, ModPred, PredHydroxy, RF-Hydroxysite) estimate prediction quality providing a confidence value. When different quality levels are provided we evaluated them as different predictors. In all figures suffix numbers in the method names indicate increasing quality threshold as defined by developers.

In addition, a random baseline method is benchmarked where predictions are obtained by randomly assigning each site as either hydroxylated or not with 50% probability. We decided not to include a separate random baseline with a probability proportional to the data imbalance since we do not really know this probability and it is difficult to estimate, in particular for hydroxylation. The random baseline represents a situation where prediction is effectively useless and predictors should achieve significantly better results to be of any practical value for experimentalists.

### 2.4 Naive HMM

The “Naive-HMM” baseline method has been implemented to demonstrate that negative examples are very different from positive sites and that they may be correctly classified by integrating new information into training datasets. A database of 750 HMMs representing hydroxylated motifs were built considering those clusters containing at least one positive site as seeds using the HMMER software (36). Naive-HMM predictions were generated by aligning all dataset sites against the HMM database. Hits with an alignment E-value better than 1.0 are considered positive predictions. The very permissive E-value is necessary because the sequence sites are very short compared to full Pfam domains. Other, less permissive, E-value thresholds do not significantly affect the performance. It should be noted that positive examples in the training and test sets overlap completely, even if negative sites inside HMM seeds are retained. The Naive-HMM baseline is therefore not meant to be of any use for biologists and does not guarantee to generalize for new sites.

### 2.5 Evaluation

The assessment is site centric, i.e. all modified (and non-modified) prolines are considered independent examples when belonging to the same protein. True positives (TP) correspond to correctly predicted hydroxylation sites whereas false positives (FP) are prolines predicted as modified in contradiction to experimental observations. True negatives (TN) are sites predicted and observed as not hydroxylated, false negatives (FN) are negative predictions of truly modified prolines. The sensitivity (Sn), specificity (Sp), weighted (or balanced) accuracy (WACC), F-measure (F1), Precision or Positive Predictive Value (PREC) and Matthew’s correlation coefficient (MCC) are computed using standard definitions (see Supplementary Material). Even where not mentioned explicitly the accuracy is always weighted.

## Acknowledgments

Fondazione Italiana per la Ricerca sul Cancro [16621] to D.P.; Associazione Italiana per la Ricerca sul Cancro [IG 17753, IG 23825] to S.T. European Union’s Horizon 2020 research and innovation programme under the Marie Skłodowska-Curie grant agreement [778247].

## Supporting Information Legends

**S1 Supplementary material.** Supplementary tables and figures.

**S2 Supplementary code and data**. Evaluation source code, predictions and reference datasets.

