## Supplementary for "Assessing predictors for new post translational modification sites: a case study on hydroxylation"

**Supplementary materials**

| **Method** | **Sn** | **Sp** | **Prec** | **Wacc** | **F1** | **Mcc** |
| --- | --- | --- | --- | --- | --- | --- |
| HydPred | 0.59 | 0.97 | 0.96 | 0.78 | 0.73 | 0.61 |
| ModPred | 0.86 | 0.78 | 0.80 | 0.82 | 0.83 | 0.64 |
| ModPred_1 | 0.78 | 0.88 | 0.87 | 0.83 | 0.82 | 0.67 |
| ModPred_2 | 0.27 | 0.99 | 0.96 | 0.63 | 0.43 | 0.38 |
| iHyd-PseAAC | 0.59 | 0.92 | 0.88 | 0.75 | 0.70 | 0.53 |
| OH-Pred | 0.78 | 0.90 | 0.88 | 0.84 | 0.83 | 0.68 |
| PredHydroxy | 0.60 | 0.98 | 0.96 | 0.79 | 0.74 | 0.63 |
| PredHydroxy_1 | 0.55 | 0.99 | 0.98 | 0.77 | 0.70 | 0.60 |
| PredHydroxy_2 | 0.46 | 0.99 | 0.98 | 0.73 | 0.63 | 0.53 |
| PredHydroxy_3 | 0.00 | 1.00 | 1.00 | 0.50 | 0.00 | 0.03 |
| RF-Hydroxysite | 0.15 | 0.27 | 0.17 | 0.21 | 0.16 | -0.59 |
| RF-Hydroxysite_1 | 0.11 | 0.41 | 0.16 | 0.26 | 0.13 | -0.50 |
| RF-Hydroxysite_2 | 0.08 | 0.60 | 0.16 | 0.34 | 0.10 | -0.38 |
| ASM3 | 0.21 | 0.82 | 0.53 | 0.51 | 0.30 | 0.03 |
| Consensus | 0.49 | 0.99 | 0.98 | 0.74 | 0.65 | 0.55 |
| Random | 0.54 | 0.50 | 0.52 | 0.52 | 0.53 | 0.03 |
| Naïve-HMM | 0.86 | 0.97 | 0.97 | 0.91 | 0.91 | 0.83 |

**Supplementary Table S1. Performance on the Literature dataset.**

| **Method** | **Sn** | **Sp** | **Prec** | **Wacc** | **F1** | **Mcc** |
| --- | --- | --- | --- | --- | --- | --- |
| HydPred | 0.08 | 0.97 | 0.75 | 0.52 | 0.14 | 0.11 |
| ModPred | 0.52 | 0.78 | 0.71 | 0.65 | 0.60 | **0.32** |
| ModPred_1 | 0.37 | 0.88 | 0.76 | 0.63 | 0.50 | 0.29 |
| ModPred_2 | 0.06 | 0.99 | 0.85 | 0.52 | 0.11 | 0.13 |
| iHyd-PseAAC | 0.08 | 0.92 | 0.50 | 0.50 | 0.14 | 0.00 |
| OH-Pred | 0.25 | 0.90 | 0.71 | 0.57 | 0.37 | 0.19 |
| PredHydroxy | 0.07 | 0.98 | 0.76 | 0.52 | 0.12 | 0.11 |
| PredHydroxy_1 | 0.02 | 0.99 | 0.64 | 0.50 | 0.03 | 0.03 |
| PredHydroxy_2 | 0.00 | 0.99 | 0.00 | 0.50 | 0.00 | -0.07 |
| PredHydroxy_3 | 0.00 | 1.00 | nan | 0.50 | 0.00 | nan |
| RF-Hydroxysite | 0.50 | 0.27 | 0.40 | 0.38 | 0.45 | -0.24 |
| RF-Hydroxysite_1 | 0.38 | 0.41 | 0.39 | 0.39 | 0.38 | -0.22 |
| RF-Hydroxysite_2 | 0.22 | 0.60 | 0.35 | 0.41 | 0.27 | -0.20 |
| ASM3 | 0.15 | 0.82 | 0.44 | 0.48 | 0.22 | -0.05 |
| Consensus | 0.02 | 0.99 | 0.66 | 0.50 | 0.04 | 0.04 |
| Random | 0.51 | 0.50 | 0.50 | 0.50 | 0.50 | 0.00 |
| Naïve-HMM | 0.87 | 0.97 | 0.97 | 0.92 | 0.91 | 0.84 |

**Supplementary Table S2. Performance on the MS-HeLa dataset.**

| **Method** | **Sn** | **Sp** | **Prec** | **Wacc** | **F1** | **Mcc** |
| --- | --- | --- | --- | --- | --- | --- |
| HydPred | 0.10 | 0.97 | 0.79 | 0.54 | 0.17 | 0.15 |
| ModPred | 0.32 | 0.78 | 0.60 | 0.55 | 0.42 | 0.12 |
| ModPred_1 | 0.22 | 0.88 | 0.65 | 0.55 | 0.33 | **0.13** |
| ModPred_2 | 0.04 | 0.99 | 0.81 | 0.52 | 0.08 | 0.10 |
| iHyd-PseAAC | 0.15 | 0.92 | 0.65 | 0.54 | 0.25 | 0.11 |
| OH-Pred | 0.17 | 0.90 | 0.63 | 0.54 | 0.27 | 0.11 |
| PredHydroxy | 0.07 | 0.98 | 0.77 | 0.53 | 0.13 | 0.12 |
| PredHydroxy_1 | 0.04 | 0.99 | 0.81 | 0.52 | 0.08 | 0.10 |
| PredHydroxy_2 | 0.03 | 0.99 | 0.76 | 0.51 | 0.05 | 0.07 |
| PredHydroxy_3 | 0.00 | 1.00 | 1.00 | 0.50 | 0.00 | 0.03 |
| RF-Hydroxysite | 0.64 | 0.27 | 0.47 | 0.46 | 0.54 | -0.10 |
| RF-Hydroxysite_1 | 0.53 | 0.41 | 0.47 | 0.47 | 0.50 | -0.06 |
| RF-Hydroxysite_2 | 0.35 | 0.60 | 0.46 | 0.47 | 0.40 | -0.06 |
| ASM3 | 0.21 | 0.82 | 0.53 | 0.51 | 0.30 | 0.03 |
| Consensus | 0.04 | 0.99 | 0.79 | 0.52 | 0.08 | 0.09 |
| Random | 0.49 | 0.50 | 0.50 | 0.50 | 0.49 | -0.01 |
| Naïve-HMM | 0.95 | 0.97 | 0.97 | 0.96 | 0.96 | 0.92 |

**Supplementary Table S3. Performance on the MS-Kim dataset.**

| **Method** | **Sn** | **Sp** | **Prec** | **Wacc** | **F1** | **Mcc** |
| --- | --- | --- | --- | --- | --- | --- |
| HydPred | 0.09 | 0.97 | 0.78 | 0.53 | 0.17 | 0.14 |
| ModPred | 0.35 | 0.78 | 0.62 | 0.57 | 0.45 | 0.15 |
| ModPred_1 | 0.24 | 0.88 | 0.67 | 0.56 | 0.36 | 0.16 |
| ModPred_2 | 0.05 | 0.99 | 0.82 | 0.52 | 0.09 | 0.11 |
| iHyd-PseAAC | 0.14 | 0.92 | 0.63 | 0.53 | 0.23 | 0.10 |
| OH-Pred | 0.19 | 0.90 | 0.65 | 0.54 | 0.29 | 0.12 |
| PredHydroxy | 0.07 | 0.98 | 0.76 | 0.52 | 0.13 | 0.12 |
| PredHydroxy_1 | 0.04 | 0.99 | 0.80 | 0.51 | 0.07 | 0.09 |
| PredHydroxy_2 | 0.02 | 0.99 | 0.72 | 0.51 | 0.04 | 0.05 |
| PredHydroxy_3 | 0.00 | 1.00 | 1.00 | 0.50 | 0.00 | 0.03 |
| RF-Hydroxysite | 0.62 | 0.27 | 0.46 | 0.44 | 0.53 | -0.12 |
| RF-Hydroxysite_1 | 0.50 | 0.41 | 0.46 | 0.46 | 0.48 | -0.09 |
| RF-Hydroxysite_2 | 0.33 | 0.60 | 0.45 | 0.46 | 0.38 | -0.08 |
| ASM3 | 0.20 | 0.82 | 0.52 | 0.51 | 0.29 | 0.02 |
| Consensus | 0.04 | 0.99 | 0.78 | 0.51 | 0.07 | 0.09 |
| Random | 0.50 | 0.50 | 0.50 | 0.50 | 0.50 | -0.01 |
| Naïve-HMM | 0.94 | 0.97 | 0.97 | 0.95 | 0.95 | 0.91 |

**Supplementary Table S4. Performance on the MS dataset (MS-Kim merged with MS-HeLa).**

| **Method** | **Sn** | **Sp** | **Prec** | **Wacc** | **F1** | **Mcc** |
| --- | --- | --- | --- | --- | --- | --- |
| HydPred | 0.80 | 0.97 | 0.97 | 0.89 | 0.88 | 0.79 |
| ModPred | 0.94 | 0.78 | 0.81 | 0.86 | 0.87 | 0.73 |
| ModPred_1 | 0.91 | 0.88 | 0.89 | 0.90 | 0.90 | 0.79 |
| ModPred_2 | 0.47 | 0.99 | 0.98 | 0.73 | 0.63 | 0.54 |
| iHyd-PseAAC | 0.80 | 0.92 | 0.91 | 0.86 | 0.85 | 0.73 |
| OH-Pred | 0.98 | 0.90 | 0.91 | 0.94 | 0.94 | 0.88 |
| PredHydroxy | 0.73 | 0.98 | 0.97 | 0.85 | 0.83 | 0.73 |
| PredHydroxy_1 | 0.64 | 0.99 | 0.99 | 0.81 | 0.77 | 0.67 |
| PredHydroxy_2 | 0.49 | 0.99 | 0.98 | 0.74 | 0.66 | 0.56 |
| PredHydroxy_3 | 0.00 | 1.00 | nan | 0.50 | 0.00 | nan |
| RF-Hydroxysite | 0.03 | 0.27 | 0.03 | 0.15 | 0.03 | -0.73 |
| RF-Hydroxysite_1 | 0.03 | 0.41 | 0.04 | 0.22 | 0.03 | -0.61 |
| RF-Hydroxysite_2 | 0.01 | 0.60 | 0.03 | 0.31 | 0.02 | -0.48 |
| ASM3 | 0.13 | 0.82 | 0.41 | 0.47 | 0.19 | -0.08 |
| Consensus | 0.64 | 0.99 | 0.98 | 0.81 | 0.77 | 0.67 |
| Random | 0.49 | 0.50 | 0.49 | 0.49 | 0.49 | -0.01 |
| Naïve-HMM | 0.87 | 0.97 | 0.97 | 0.92 | 0.91 | 0.84 |

**Supplementary Table S5. Performance on the Literature-collagen dataset.**

| **Method** | **Sn** | **Sp** | **Prec** | **Wacc** | **F1** | **Mcc** |
| --- | --- | --- | --- | --- | --- | --- |
| HydPred | 0.23 | 0.97 | 0.90 | 0.60 | 0.37 | 0.30 |
| ModPred | 0.85 | 0.78 | 0.80 | 0.82 | 0.82 | 0.64 |
| ModPred_1 | 0.77 | 0.88 | 0.87 | 0.82 | 0.81 | 0.65 |
| ModPred_2 | 0.22 | 0.99 | 0.96 | 0.61 | 0.36 | 0.33 |
| iHyd-PseAAC | 0.44 | 0.92 | 0.85 | 0.68 | 0.58 | 0.41 |
| OH-Pred | 0.69 | 0.90 | 0.87 | 0.79 | 0.77 | 0.60 |
| PredHydroxy | 0.41 | 0.98 | 0.95 | 0.69 | 0.57 | 0.47 |
| PredHydroxy_1 | 0.26 | 0.99 | 0.96 | 0.63 | 0.41 | 0.37 |
| PredHydroxy_2 | 0.10 | 0.99 | 0.93 | 0.55 | 0.19 | 0.21 |
| PredHydroxy_3 | 0.01 | 1.00 | 1.00 | 0.51 | 0.02 | 0.09 |
| RF-Hydroxysite | 0.19 | 0.27 | 0.21 | 0.23 | 0.20 | -0.55 |
| RF-Hydroxysite_1 | 0.14 | 0.41 | 0.19 | 0.27 | 0.16 | -0.47 |
| RF-Hydroxysite_2 | 0.08 | 0.60 | 0.17 | 0.34 | 0.11 | -0.37 |
| ASM3 | 0.08 | 0.82 | 0.32 | 0.45 | 0.13 | -0.14 |
| Consensus | 0.17 | 0.99 | 0.94 | 0.58 | 0.28 | 0.27 |
| Random | 0.49 | 0.50 | 0.49 | 0.49 | 0.49 | -0.02 |
| Naïve-HMM | 0.75 | 0.97 | 0.96 | 0.86 | 0.84 | 0.73 |

**Supplementary Table S6. Performance on the MS-collagen dataset.**

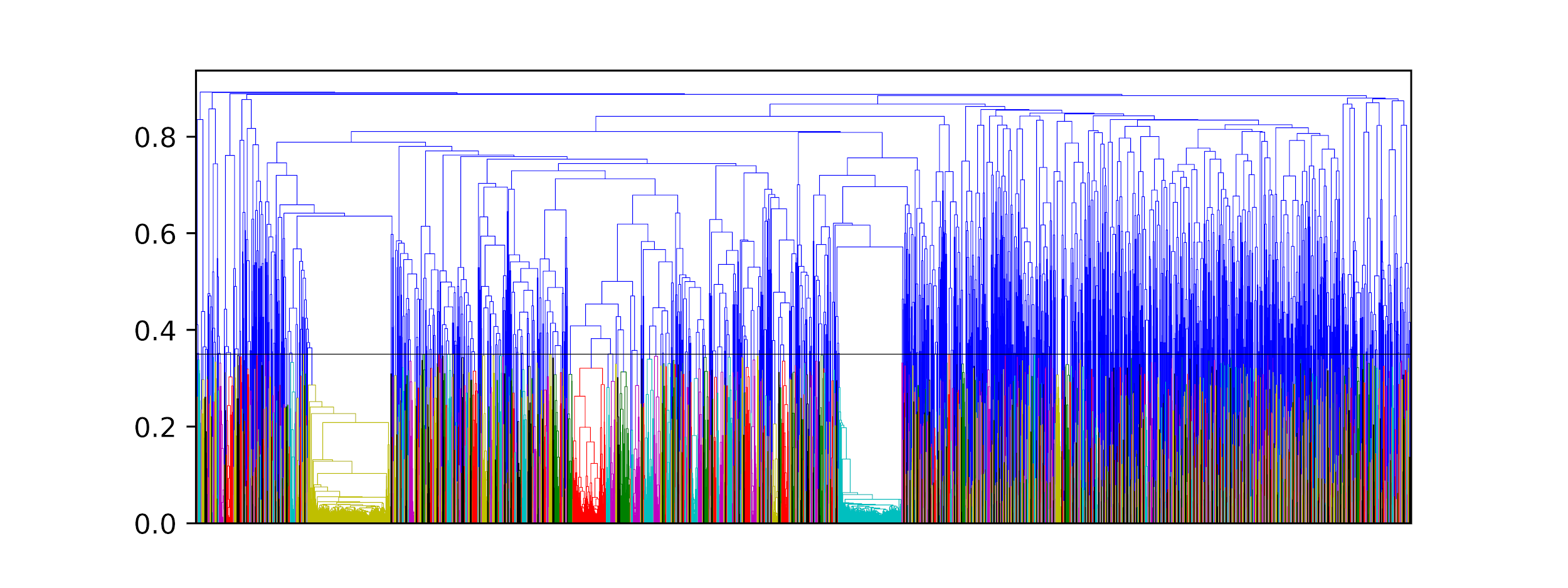

**Supplementary Figure S1. Dendrogram of hydroxylated and non hydroxylated sites.**

Nodes of the same color have a UPGMA distance closer than 0.35. The distance is calculated as the inverse of the sum of the Blosum62 score for each pair of residues with a penalty of -5 and -1 for gap opening and extension respectively (gaps are only possible for sites shorter than the window, i.e. close to the sequence end). The three larger groups (yellow, red, ciano) correspond to three different collagen motifs.

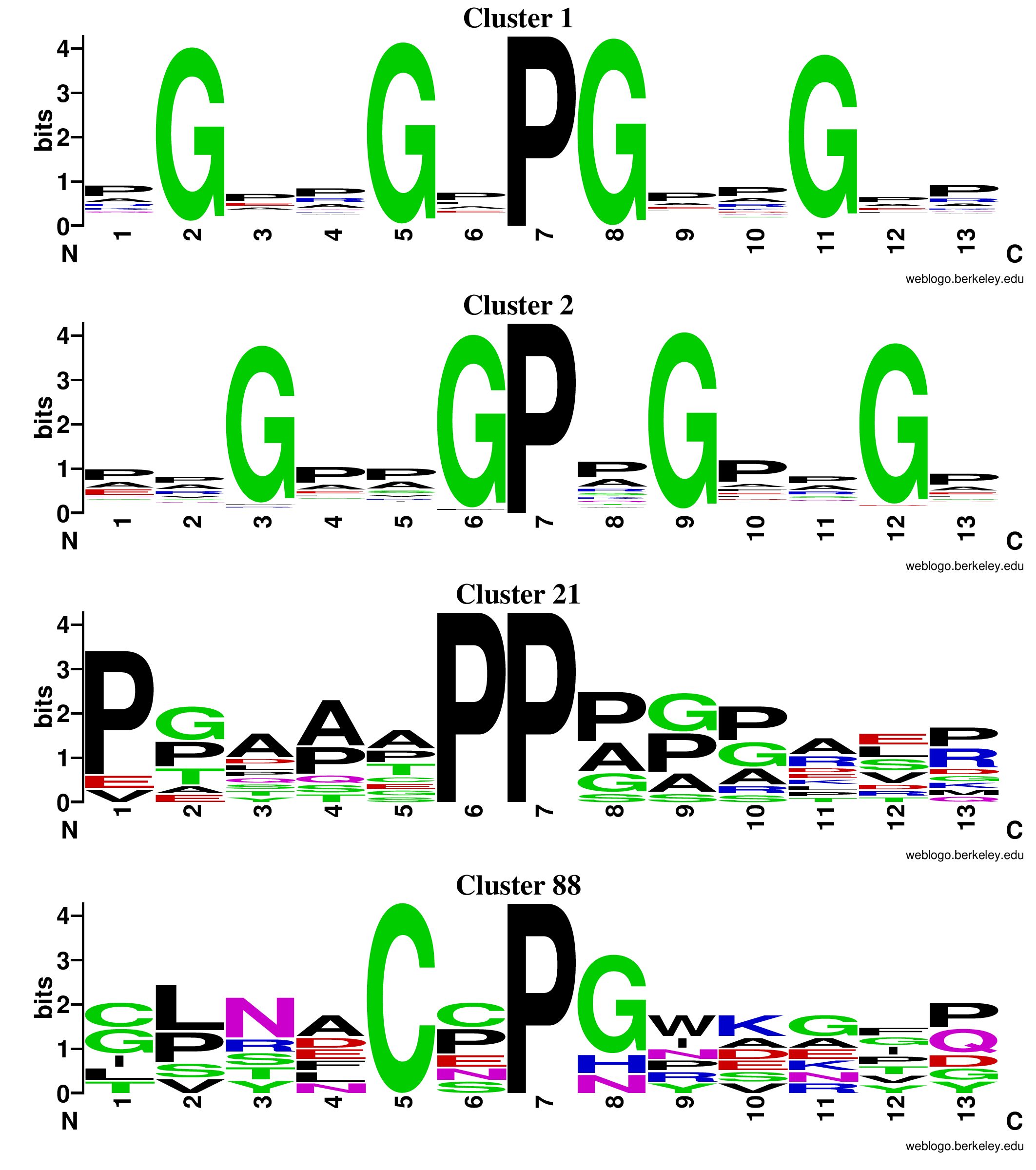

**Supplementary Figure S2. Sequence logo of collagen clusters**

The fifth logo mentioned in the manuscript is missing as the corresponding cluster, #559, contains only one site sequence, “CSFECQPARGPPG”.

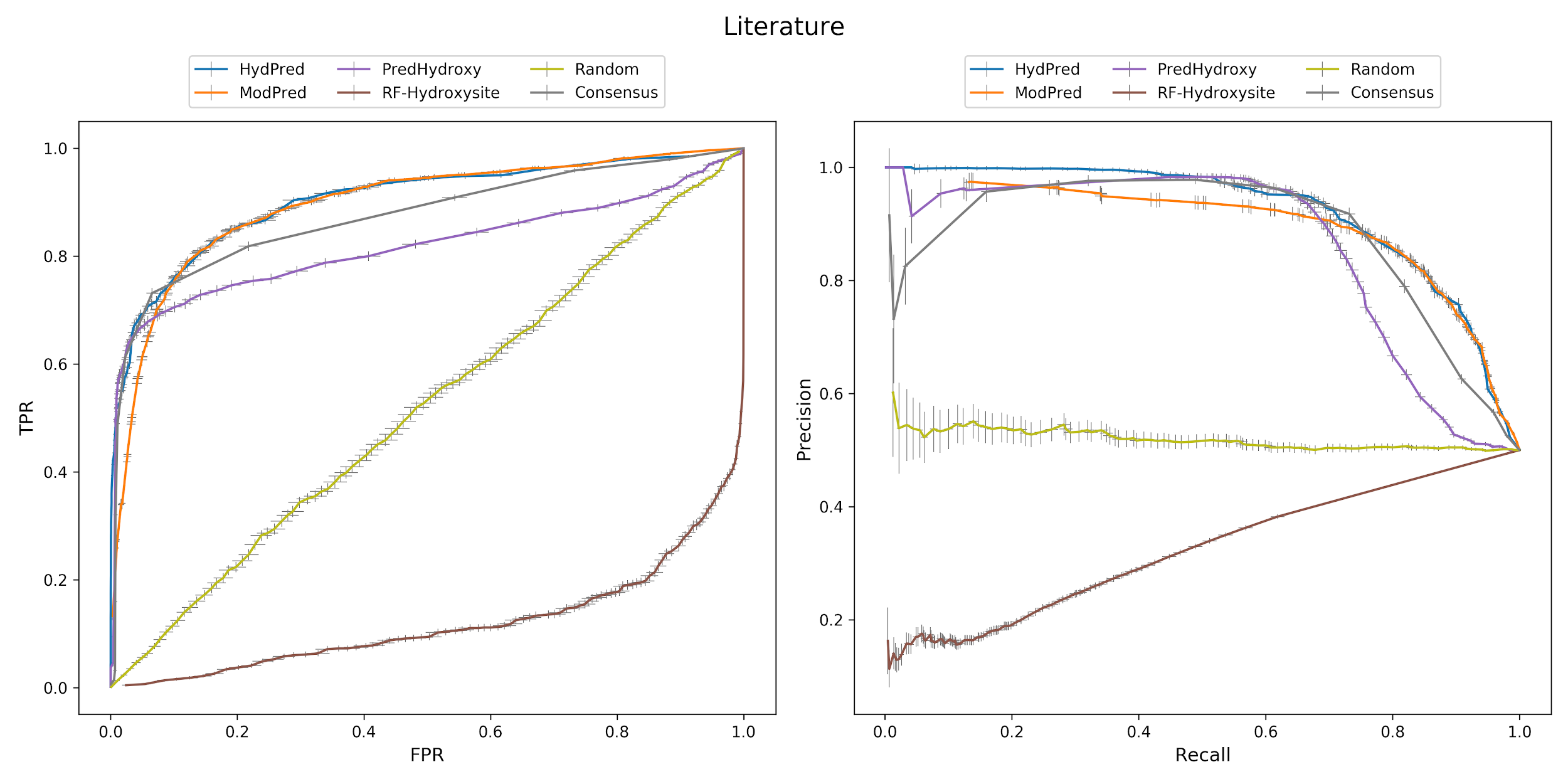

**Supplementary Figure S3. ROC and precision-recall curves on the Literature dataset.**

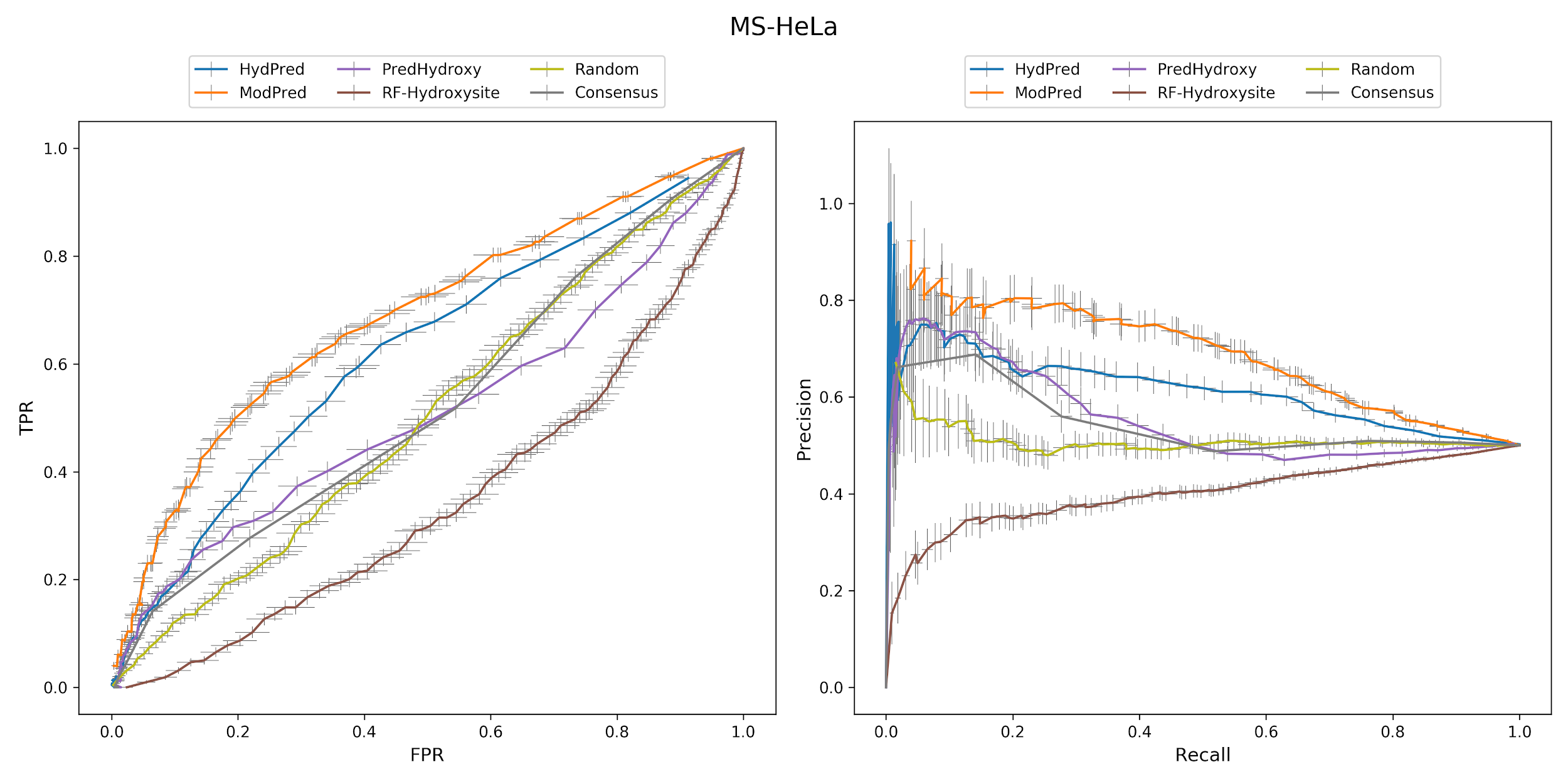

**Supplementary Figure S4. ROC and precision-recall curves on the MS-HeLa dataset.**

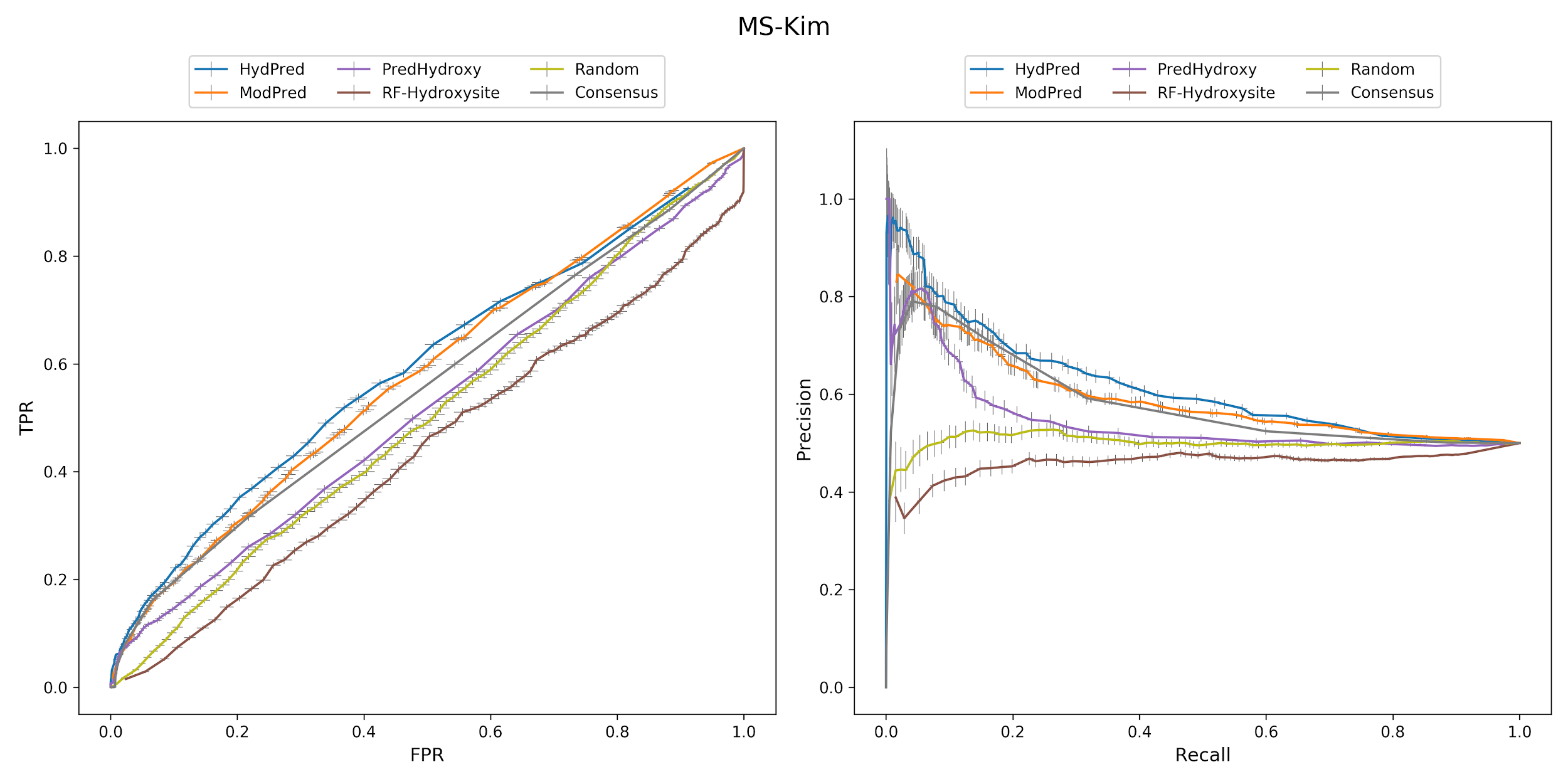

**Supplementary Figure S5. ROC and precision-recall curves on the MS-Kim dataset.**

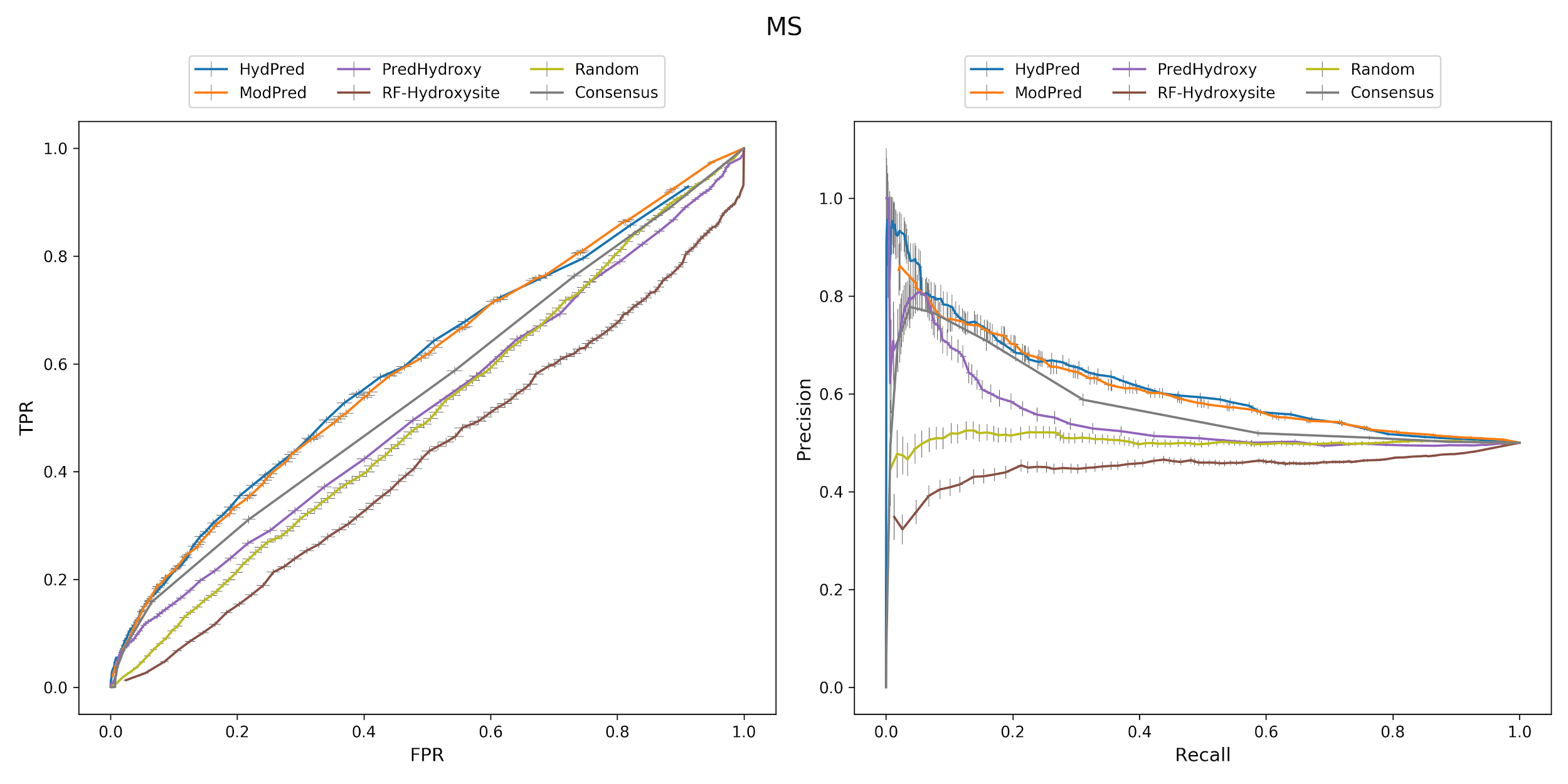

**Supplementary Figure S6. ROC and precision-recall curves on the MS dataset (MS-Kim merged with MS-HeLa).**

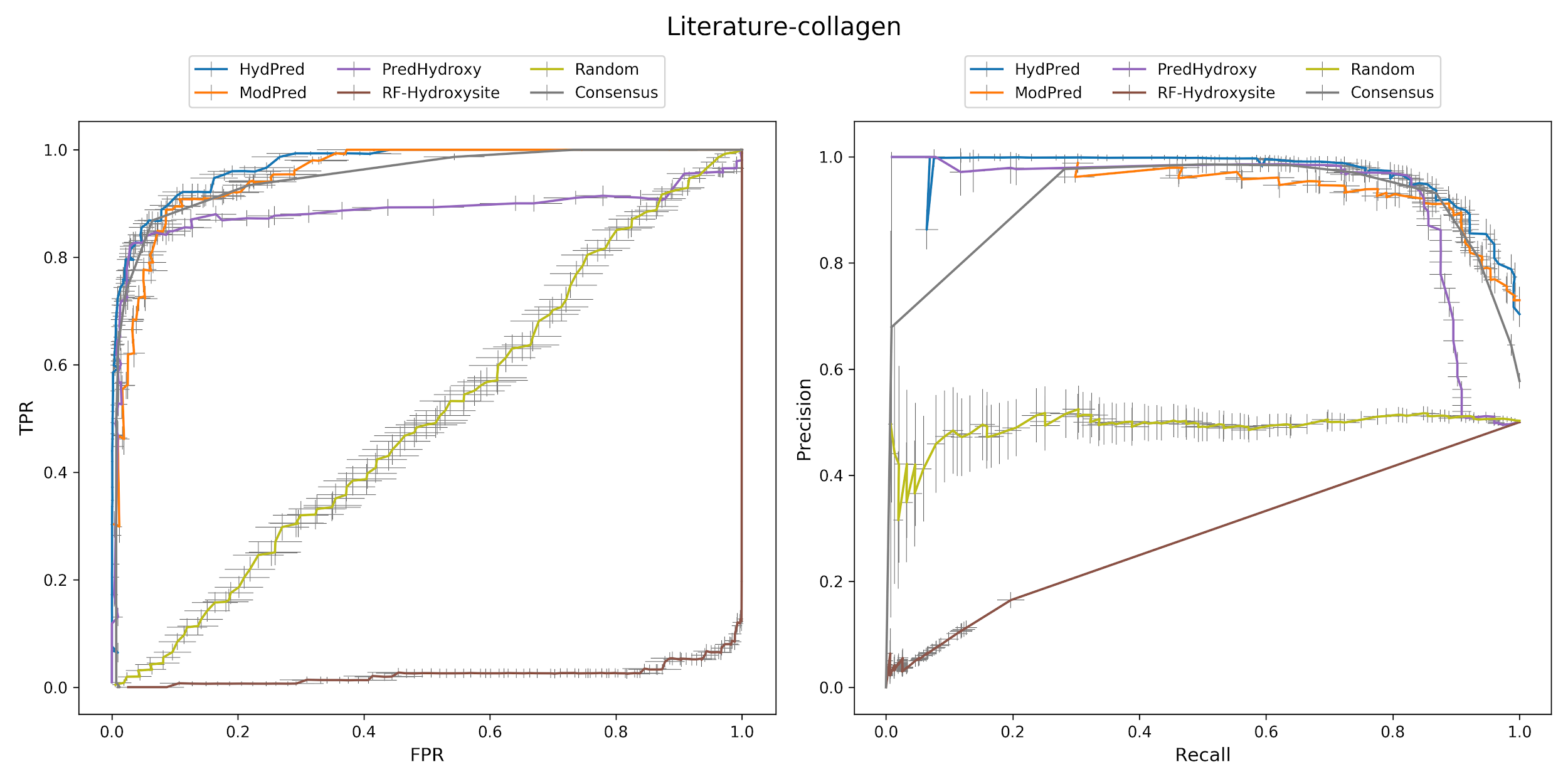

**Supplementary Figure S7. ROC and precision-recall curves on the Literature-collagen dataset.**

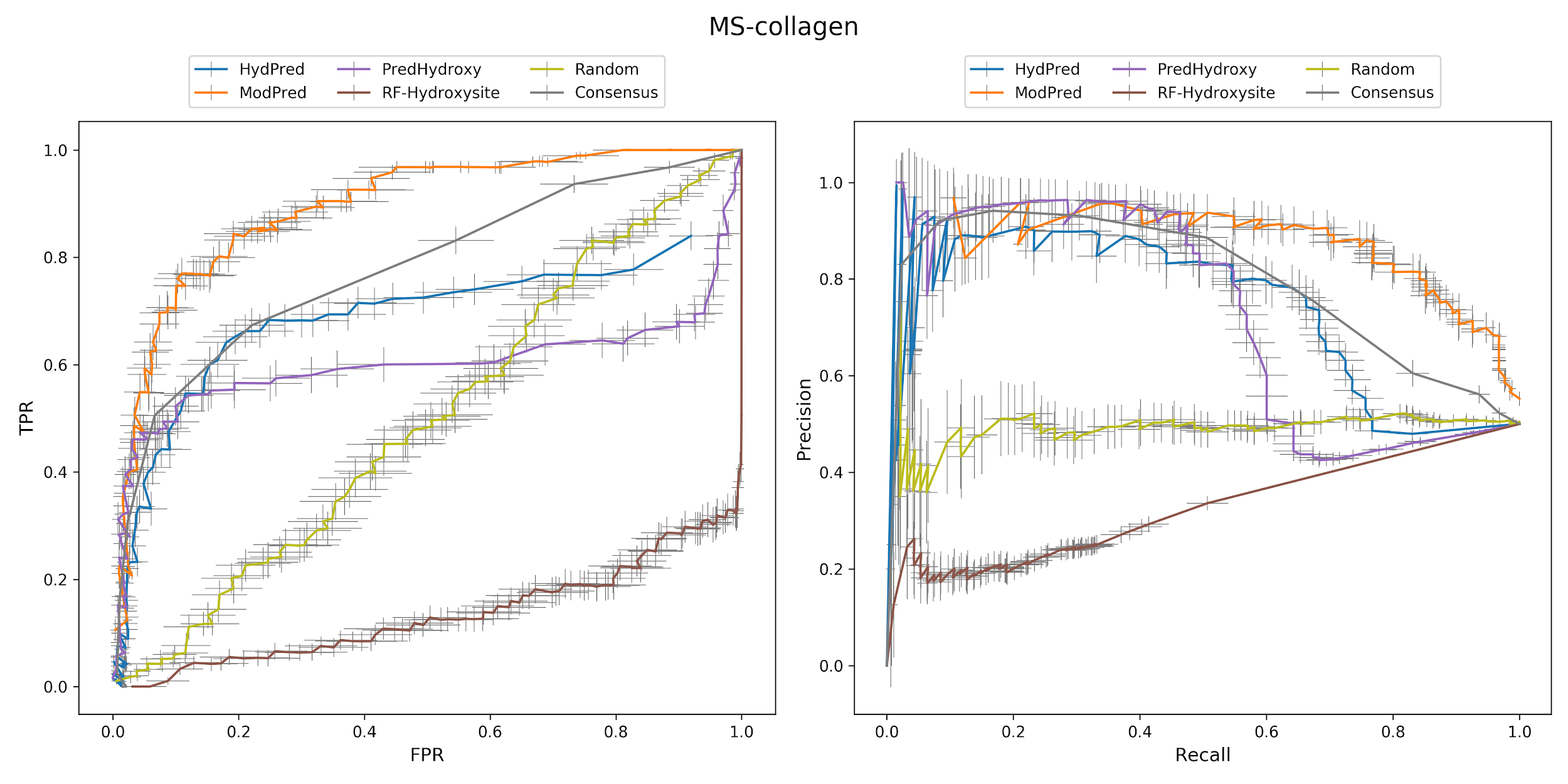

**Supplementary Figure S8. ROC and precision-recall curves on the MS-collagen dataset.**
